## Supplementary Figures for "Convergent evolution between PALI1 and JARID2 for the allosteric activation of PRC2"

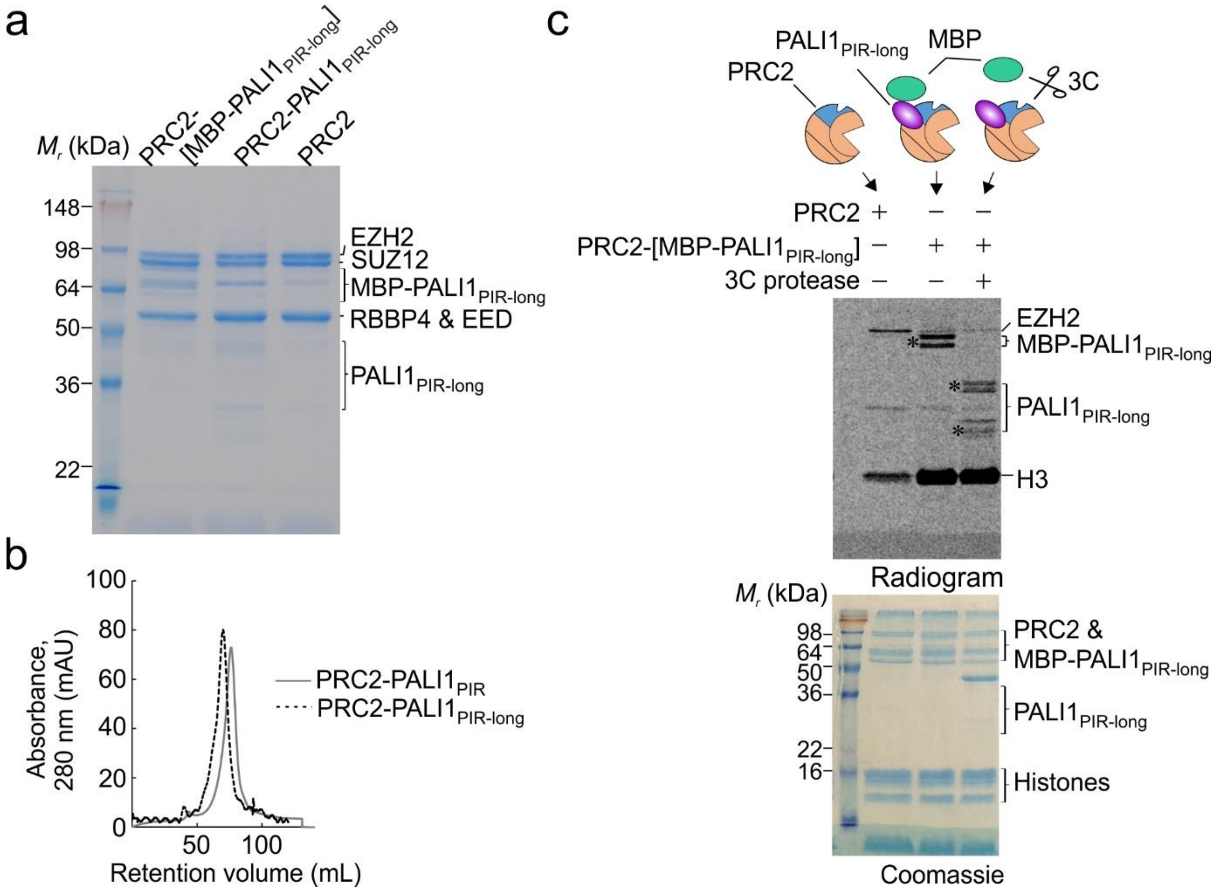


**Supplementary Fig. 1. PALI1 is methylated *in vitro*.**

**a,** Coomassie blue-stained SDS-PAGE of recombinant human PRC2-PALI1$_{PIR-long}$ complexes, as indicated. **b,** Gel filtration chromatography of the PRC2-PALI1$_{PIR}$ and PALI1$_{PIR-long}$ complexes (HiPrep 16/600 Sephacryl S-400 HR column). **c,** HTMase assay of the PRC2-[MBP-PALI1$_{PIR-long}$] complex with a mononucleosome substrate performed in the presence or absence of 3C protease to confirm that PALI1$_{PIR-long}$ is methylated. The MBP-cleaved and uncleaved PALI1$_{PIR-long}$ band indicated on the radiogram with asterisks

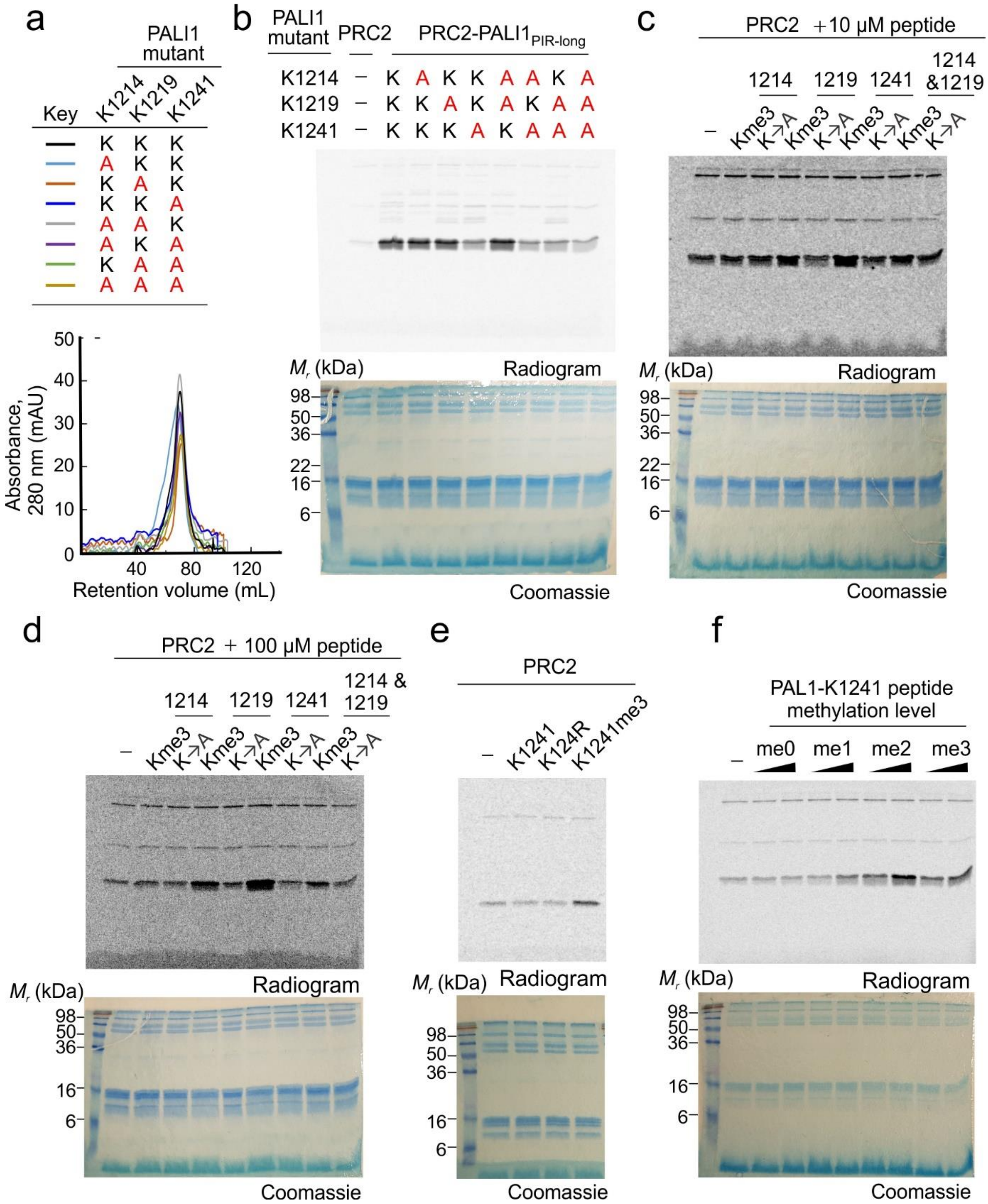


**Supplementary Fig. 2. PALI1-K1241me2/3 is required and sufficient to stimulate the HMTase activity of PRC2.**

**a,** Gel filtration chromatography (HiPrep 16/600 Sephacryl S-400 HR column) of the PRC2-PALI1$_{PIR\text{-}long}$ wild type and mutants, as indicated. **b,** A representative full radiogram and the corresponding uncropped Coomassie blue-stained SDS-PAGE, as shown in Fig. 2a. **c,** A representative full radiogram and the corresponding uncropped Coomassie blue-stained SDS-PAGE as shown in Fig. 2b for 10 µM peptide concentration. **d,** A representative full radiogram and the corresponding uncropped Coomassie blue-stained SDS-PAGE as shown in Fig. 2b for 100 µM peptide concentration. **e,** A representative full radiogram and the corresponding uncropped Coomassie blue-stained SDS-PAGE as shown in Fig. 2c. **f,** A representative full radiogram and the corresponding uncropped Coomassie blue-stained SDS-PAGE as shown in Fig. 2d.

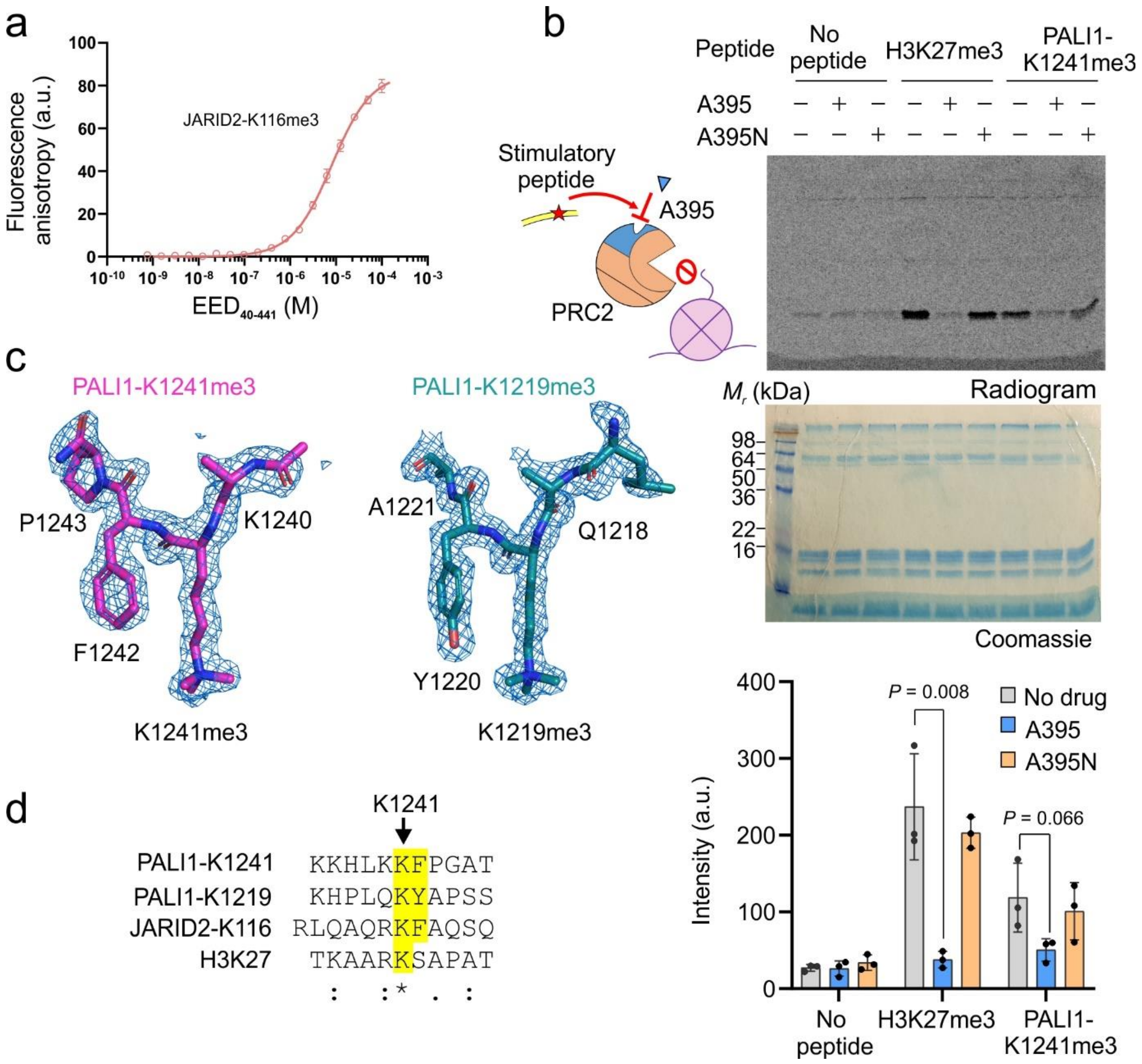


**Supplementary Fig. 3. PALI1-K1241me2/3 binds to the aromatic cage of the regulatory subunit EED to stimulate PRC2.**

**a,** Fluorescence anisotropy performed to quantify the binding affinity of EED for the 5-FAM labelled JARID2-K116me3 peptide. Error bars represent standard deviation over three independent experiments carried out in different days. Dissociation constants ($K_d$) and 95% confidence bounds on the coefficient are indicated in Fig. 3a. **b,** HMTase assay of 200 nM PRC2 using 2 μM mononucleosomes, in the presence or absence of 50 μM stimulatory peptides, as indicated, and in the presence or absence of 0.8 μM allosteric inhibitor A395 or the negative control compound A395N. The bar plot represents the means of quantification using densitometry from three independent replicates with the error bars represent standard deviation. P-values were determined using unpaired two-tailed Student's t-test. **c,** Fo-Fc omit electron density maps for PALI1 peptides bound to EED, contoured at 3.0 σ. Omit map was calculated using Polder Maps[65] in PHENIX[62] visualized using PyMOL (The PyMOL Molecular Graphics System, Version 2.0 Schrödinger, LLC.). **d,** The sequences of the tri-methyl-lysine peptides used for the crystallization of EED, including the PALI1 peptides (this study), the JARID-K116me3 peptide[21] and the H3K27me3 peptide[23], aligned according to the methylated lysine residues. The methyl-lysine and the adjacent aromatic amino acids at the +1 position are highlighted in each of the peptides, when applicable.

a

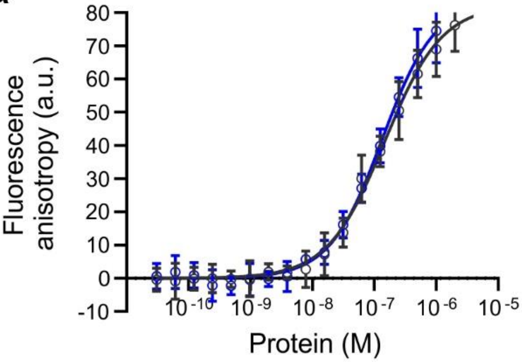


| Key | Protein | Probe | $K_d$ (nM) | Hill |
|---|---|---|---|---|
| (gray) | PRC2 | G-tract RNA | 147 ± 22 | 0.94 ± 0.09 |
| (blue) | PRC2-PALI1$_{PIR}$ | G-tract RNA | 134 ± 25 | 1.02± 0.11 |

b

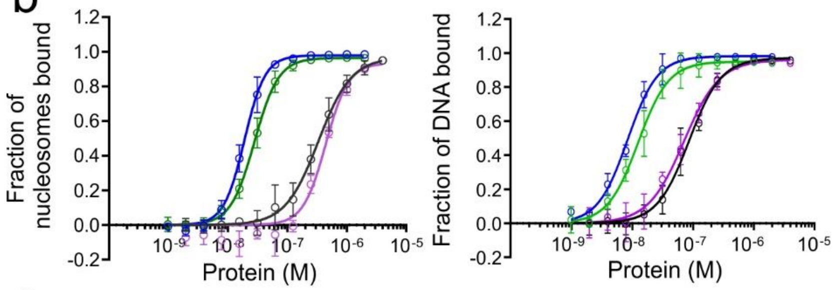


c

| Key | Protein | Nucleosome binding | | DNA binding | |
|---|---|---|---|---|---|
| | | $K_d$(nM) | Hill | $K_d$(nM) | Hill |
| (gray) | PRC2 | 333 ± 30 | 1.56 ± 0.17 | 86.2 ± 5.5 | 1.65 ± 0.14 |
| (purple) | PRC2 + 100 μM PALI1-K1241me3 peptide | 444 ± 37 | 2.27 ± 0.36 | 70.6 ± 5.6 | 1.48 ± 0.14 |
| (blue) | PRC2-PALI1$_{PIR}$ | 19.0 ± 0.6 | 2.49 ± 0.18 | 8.35 ± 0.34 | 1.76 ± 0.11 |
| (green) | PRC2-PALI1$_{PIR}$ K1241A | 28.1 ± 1.0 | 2.15 ± 0.14 | 12.5 ± 0.8 | 1.73 ± 0.17 |

**Supplementary Fig. 4. PALI1 is a DNA binding subunit of PRC2.**

**a,** Fluorescence anisotropy was performed to quantify the affinity of PRC2 complexes for G4 24 RNA $(UUAGGG)_4$. Data represent the mean of three independent experiments that were carried out on different days. Error bars represent standard deviation. Standard errors of dissociation constants ($K_d$) and Hill coefficients are indicated in the table. **b, c,** Quantification of EMSA from Fig. 4b. The affinities of the indicated PRC2 complexes to mononucleosomes and free-DNA, from data shown in Fig. 4b, were quantified using densitometry from the free-nucleosome and free-DNA bands. Data represent the mean of three independent experiments and the error bars represent standard deviations. Standard errors of dissociation constants ($K_d$) and Hill coefficients are indicated in the table.

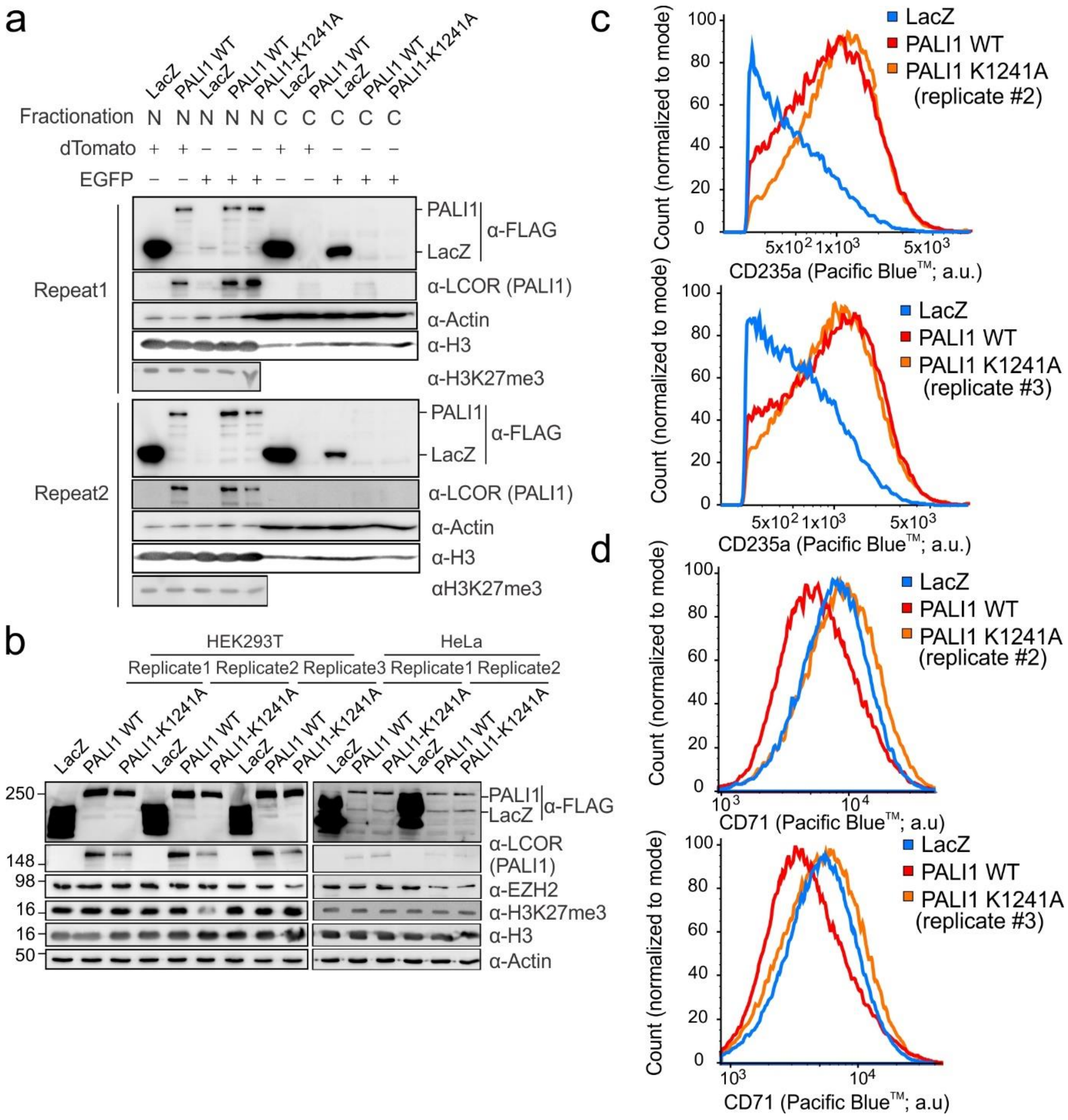


**Supplementary Fig. 5. Overexpression of PALI1 reduces K562 cell proliferation, with the effect alleviated by K1241A**

**a,** Nuclear and cytoplasmic fractions of K562 cells overexpressing different proteins were isolated and examined by western blotting with antibodies as indicated. Results are from two independent replicates are shown. Subcellular fractions are abbreviated as N for nuclear fraction and C for cytoplasmic fraction. **b,** Western blots of HEK293T and HeLa cells overexpressing proteins as indicated performed using antibodies as indicated. Results from two or three independent replicates are shown. **c, d,** The two replicates of the histograms representing the distribution of cells based on the expression of the erythroid marker CD235a (c) or CD71 (d) as detected by flow cytometric analysis of K562 cells overexpressing different proteins, as indicated (Blue: LacZ, red: PALI1 WT, orange: PALI1 K1241A). Results are from three independent experiments that were carried out on different days, starting each time from lentiviral transduction, with the other replicate presented in a body figure (Fig 5d).

**SUPPLEMENTARY MATERIAL**

**Supplementary Table 1. The summary of the PRC2 methylome *in vivo* and *in vitro*.** Residues with a position probability of less than 0.95 were indicated with red text and probability scores shown in parentheses. Residues from peptides that were ambiguous between EZH1 and EZH2 are indicated by an asterisk.

**Supplementary Table 2. Primers and sequences used in this study.**

**Supplementary Notes. MS/MS spectra of the methylated PALI1 K1241 as identified *in vivo* and *in vitro*.**
