## Supplementary Table 2 for "Convergent evolution between PALI1 and JARID2 for the allosteric activation of PRC2"

**Supplementary Table 2: Primers and sequences.**

**Cloning Primers**

PALI1_F2_pFB1.HMBP.PrS GGAAGTTCTGTTCCAGGGGCCCGGGCAGCGAATGATCCAACAATTTG

PALI1_R1557_pFB1.HMBP.PrS
ATGCCTCGAGACTGCAGGCTCTAGATTATCACTTTGCATCCAGCCGCCTCCG

PALI1_R310_inter GACTTTCTACTAAAGCTGAACTCTGCATATGGTCTTTACCATCCTCACA

PALI1_F312_inter CATATGTGAGGATGGTAAAGACCATATGCAGAGTTCAGCTTTAGTAGAA

PALI1_F1058_pFB1.HMBP.PrS GGAAGTTCTGTTCCAGGGGCCCGGGACTTCAGAAAAGGAAGCTGC

PALI1_R1329_pFB1.HMBP.PrS
ATGCCTCGAGACTGCAGGCTCTAGATTATCAATTCTTGATTAAACGTTGCTG

PALI1_R1250_pFB1.HMBP.PrS
ATGCCTCGAGACTGCAGGCTCTAGATTATCAATTCTTAGCAGGGGTAGCTCC

EED_F40_pGEX-MHL ttgtatttccagggcGACGCTGTCAGTATAGAAAGTG

EED_F76_pGEX-MHL ttgtatttccagggcAAGAAATGCAAATATTCTTTCAAATG

EED_R441_pGEX-MHL caagcttcgtcatcaTCGAAGTCGATCCCAGCGC

PALI1_F1_pHIV-EGFP/pHIV-dTomato
AACTATTCTAGAGTACCCACCATGGACTACAAAGACGATGACGACAAGATGCAGCGAATGATCCAAC
AA

PALI1_R1557_pHIV-EGFP/pHIV-dTomato
AGGGGCGGATCCTAGCCCCTATTATACCTTTCTCTTCTTTTTTGGCTTTGCATCCAGCCGCCTCCG

LacZ_F_pHIV-EGFP/ pHIV-dTomato
AACTATTCTAGAGTACCCACCATGGACTACAAAGACGATGACGACAAGGTCGTTTTACAACGTCGTG
AC

LacZ_R_pHIV-EGFP/pHIV-dTomato
AGGGGCGGATCCTAGCCCCTATTATACCTTTCTCTTCTTTTTTGGTTTTTGACACCAGACCAACTG

**Primers used to generate point mutations**

PALI1_K1241A_F TTGAAGGCTTTTCCTGGAGCTACCCCT

PALI1_K1241A_R AGGAAAAGCCTTCAAGTGCTTTTTCAA

PALI1_K1214A_F CCTGTCGCTCATCCTCTTCAGAAATAC

PALI1_K1214A_R AGGATGAGCGACAGGGGGAACGTCTCC

PALI1_K1219A_F CTTCAGGCTTACGCTCCTTCCAGCCTA

PALI1_K1219A_R AGCGTAAGCCTGAAGAGGATGCTTGAC

PALI1_K1214A/K1219A_R AGCGTAAGCCTGAAGAGGATGAGCGAC

**PALI1 full-length senqeunce**

ATGCAGCGAATGATCCAACAATTTGCTGCTGAATATACCTCAAAAAATAGCTCTACTCAGGACCCCAGCCAGC
CCAATAGCACAAAGAACCAAAGCCTGCCGAAAGCATCTCCAGTCACCACCTCTCCCACGGCTGCAACTACTCA
GAACCCTGTGCTCAGCAAACTTCTCATGGCTGACCAAGACTCACCTCTGGACCTTACTGTCAGAAAGTCTCAGT
CAGAACCTAGCGAACAAGACGGTGTACTTGATCTGTCCACTAAGAAAAGTCCATGTGCTGGCAGCACTTCCCT
GAGCCACTCTCCAGGCTGCTCCAGTACTCAAGGGAACGGTGAGAACTCAACAGAGGCAAAAGCAGTAGATTC
TAACAATCAGTCGAAGTCCCCACTGGAGAAATTTATGGTCAAACTGTGTACTCATCATCAAAAGCAATTCATTC
GTGTTCTGAACGACCTGTACACTGAATCTCAACCAGGCACTGAGGACCTGCAGCCTTCTGATTCGGGAGCAAT
GGATGTATCCACTTGCAATGCTGGCTGTGCCCAGCTCAGCACCAAACATAAGGAAAAAGATGCTCTGTGTCTC
GATATGAAGTCTTCTGCTTCTGTAGATTTGTTCGTAGACTCGTCAGACTCTCACAGCCCTCTACACTTGACGGA
ACAGACCCCGAAGAAGCCTCCTCCTGAGATAAACCCTGTAGATGGAAGAGAGAATGCCTTGACTGTTGTCCA
GAAAGATTCCTCTGAACTTCCAACCACTAAATCGAATTCTATTAATAGCAGTTCAGTGGATAGTTTCACTCCGG
GATACCTCACTGCATCTAATTGTTCCTCAGTGAACTTCCACCACATCCCTAAAATCTTGGAGGGGCAGACCACT
GGACAAGAGCAAGACACAAATGTGAACATATGTGAGGATGGTAAAGACCATATGCAGAGTTCAGCTTTAGTA
GAAAGTCTAATTACAGTAAAAATGGCAGCTGAGAATAGTGAGGAAGGCAATACCTGTATTATTCCTCAAAGA
AATTTGTTCAAAGCTTTATCAGAAGAGGCTTGGAACTCAGGGTTTATGGGGAACTCATCTAGAACTGCTGACA
AAGAGAATACTTTACAGTGTCCAAAAACACCTTTGCGCCAGGATTTAGAGGCAAATGAACAAGATGCAAGGC
CAAAGCAAGAGAACCATCTTCACTCTCTGGGAAGAAATAAGGTGGGTTACCATTTACATCCCAGTGATAAGGG
CCAGTTTGATCATTCCAAAGATGGTTGGTTAGGCCCCGGCCCTATGCCAGCTGTACACAAAGCGGCAAATGGA
CACTCAAGAACCAAGATGATATCAACCTCCATCAAGACAGCTCGGAAAAGTAAAAGGGCATCAGGGCTGAGG
ATAAATGATTATGATAACCAGTGTGATGTTGTTTATATCAGTCAACCAATAACAGAATGCCACTTTGAGAATCA
AAAATCAATATTATCTTCTCGGAAAACAGCCAGAAAGAGTACTCGAGGATACTTTTTCAATGGTGACTGTTGT
GAGCTGCCAACTGTTCGTACACTGGCCAGAAATTTACACTCCCAGGAAAAAGCAAGCTGCTCAGCATTGGCAT
CAGAGGCAGTTTTCACTCCTAAGCAGACCCTTACAATTCCAGCCCCTAGACATACAGTAGATGTGCAGCTTCCC
AGAGAAGACAACCCTGAAGAACCTAGCAAGGAAATCACCTCTCACGAGGAAGGAGGTGGAGACGTTTCACCT
CGAAAAGAACCTCAAGAGCCTGAGGTTTGCCCCACAAAGATTAAGCCGAACCTGAGCAGCTCCCCTAGGTCA
GAGGAAACGACAGCCTCCAGCCTGGTGTGGCCTCTCCCTGCTCACCTTCCTGAAGAGGACCTGCCAGAAGGT
GGCTCCACAGTCTCAGCTCCCACAGCAAGTGGGATGTCTTCTCCTGAACACAACCAACCACCAGTTGCACTGTT
GGATACGGAGGAGATGAGTGTACCCCAGGACTGTCACCTCCTTCCCTCCACTGAAAGCTTTTCCGGGGGAGTC
AGTGAAGATGTCATTTCTAGGCCTCATTCTCCTCCTGAAATAGTCAGTAGAGAAGAAAGTCCTCAGTGCTCAG
AAAATCAGAGTTCCCCAATGGGCTTGGAGCCCCCCATGAGTCTGGGAAAGGCTGAGGACAACCAAAGCATCA
GTGCTGAGGTTGAGTCTGGAGACACCCAGGAGCTAAATGTCGACCCACTCTTGAAGGAAAGCAGCACTTTTA
CTGATGAAAACCCCAGTGAAACTGAGGAAAGTGAGGCAGCAGGTGGTATAGGAAAATTAGAGGGAGAGGA
CGGTGATGTAAAATGCCTGTCAGAAAAAGACACGTATGATACAAGCATTGACTCACTCGAAGAGAATTTGGA
CAAGAAGAAAAAAGGTAAAAAATTCCCTGAGGCCTCTGATAGGTGCCTAAGAAGTCAACTTTCGGATTCTTCC
TCTGCTGACAGATGCCTAAGAAATCAGAGTTCAGATTCTTCCTCAGCTTGTCTTGAAATCAAAGTTCCTAAAAA
TCCTAGTGCAAAACGTTCAAAAAAAGAAGGGCACCCTGGTGGGACAACACCTAAGGGCCTTCTACCTGACAG
TTTCCACACGGAAACTCTGGAGGACACAGAAAAGCCAAGTGTCAATGAACGCCCCTCTGAGAAAGATGCTGA
GCAGGAGGGCGAAGGCGGGGGGATCATCACCAGGCAGACTTTGAAAAACATGCTGGACAAAGAAGTCAAG
GAGTTACGAGGAGAGATTTTCCCCAGCAGGGACCCCATAACCACAGCTGGACAGCCACTGCCTGGAGAGAGA
TTGGAAATCTATGTTCAGTCTAAAATGGATGAGAAGAATGCTCATATCCCCTCAGAAAGTATTGCTTGTAAGA
GGGACCCAGAACAGGCAAAAGAAGAGCCAGGGCATATTCCCACACAGCATGTGGAGGAGGCTGTGAATGAG
GTAGACAACGAAAACACCCAGCAGAAAGATGATGAGAGTGATGCCCCATGCAGCTCTCTTGGGTTGTCGAGT
AGTGGAAGTGGTGATGCTGCTAGGGCACCAAAATCGGTGCCAAGGCCTAAAAGATTGACCTCTTCAACCTAC
AACCTAAGACACGCTCATTCTCTGGGCTCCTTGGATGCTTCAAAAGTGACTTCAGAAAAGGAAGCTGCACAAG
TAAACCCCATAATGCCAAAGGAAAATGGAGCTTCAGAGAGTGGAGACCCCCTAGATGAGGACGATGTTGACA
CCGTGGTAGATGAACAGCCAAAGTTTATGGAATGGTGTGCTGAGGAGGAGAACCAAGAGCTCATCGCCAACT
TCAATGCCCAGTACATGAAAGTTCAGAAGGGCTGGATCCAGTTGGAGAAAGAAGGACAGCCAACACCAAGA
GCAAGGAACAAATCAGATAAACTGAAAGAGATTTGGAAAAGCAAGAAAAGGTCACGGAAATGTAGGAGTTC
ATTGGAGAGTCAGAAGTGTTCTCCTGTTCAGATGCTCTTTATGACAAACTTTAAATTATCTAATGTTTGTAAATG
GTTCTTAGAGACAACTGAAACCCGGTCTCTAGTCATTGTGAAGAAGCTCAATACTCGCCTTCCAGGAGACGTT

CCCCCTGTCAAGCATCCTCTTCAGAAATACGCTCCTTCCAGCCTATATCCCAGTTCACTACAGGCTGAGCGCTTG
AAAAAGCACTTGAAGAAATTTCCTGGAGCTACCCCTGCTAAGAATAATTGGAAAATGCAGAAGCTCTGGGCCA
AATTTCGAGAGAATCCTGATCAAGTGGAGCCAGAAGATGGCAGTGATGTCAGCCCCGGCCCTAATTCTGAAG
ACAGCATAGAGGAAGTCAAGGAAGATAGAAACAGTCATCCTCCAGCAAACCTGCCCACTCCAGCCAGTACCC
GGATTCTTAGAAAATATTCCAATATTCGAGGAAAGCTCAGAGCCCAGCAACGTTTAATCAAGAATGAGAAAAT
GGAATGCCCAGATGCTCTGGCTGTGGAAAGTAAGCCAAGTCGTAAGAGCGTATGCATCAACCCTCTGATGTCC
CCCAAGCTTGCCCTGCAAGTGGATGCAGATGGGTTTCCTGTTAAGCCCAAGAGTACTGAAGGAATGAAGGGA
AGGAAGGGGAAGCAGGTGTCTGAAATCTTGCCTAAAGCAGAAGTTCAGAGTAAACGCAAGAGAACAGAAGG
CAGCAGCCCTCCAGATAGTAAGAACAAGGGGCCTACGGTGAAAGCCAGCAAAGAAAAGCATGCTGATGGAG
CCACCAAAACCCCTGCTGCCAAGAGGCCAGCTGCAAGGGACAGAAGCAGCCAACCCCCCAAAAAGACGTCTT
TGAAAGAGAATAAAGTGAAGATCCCTAAAAAGTCCGCTGGGAAGAGCTGCCCTCCCTCCAGGAAAGAAAAAG
AGAATACAAACAAAAGGCCTTCCCAGTCTATTGCCTCGGAAACACTGACGAAACCTGCAAAACAGAAGGGGG
CCGGTGAATCCTCTTCAAGGCCTCAGAAAGCCACGAATAGGAAGCAGAGTAGTGGAAAGACTCGGGCCAGAC
CCTCAACGAAAACCCCAGAGAGCAGTGCAGCTCAGAGAAAGCGAAAGCTGAAGGCAAAGCTGGACTGTTCG
CACAGCAAACGGAGGCGGCTGGATGCAAAG
