## Supplementary Notes for "Convergent evolution between PALI1 and JARID2 for the allosteric activation of PRC2"

PALI1 K1241me2

*in vitro*

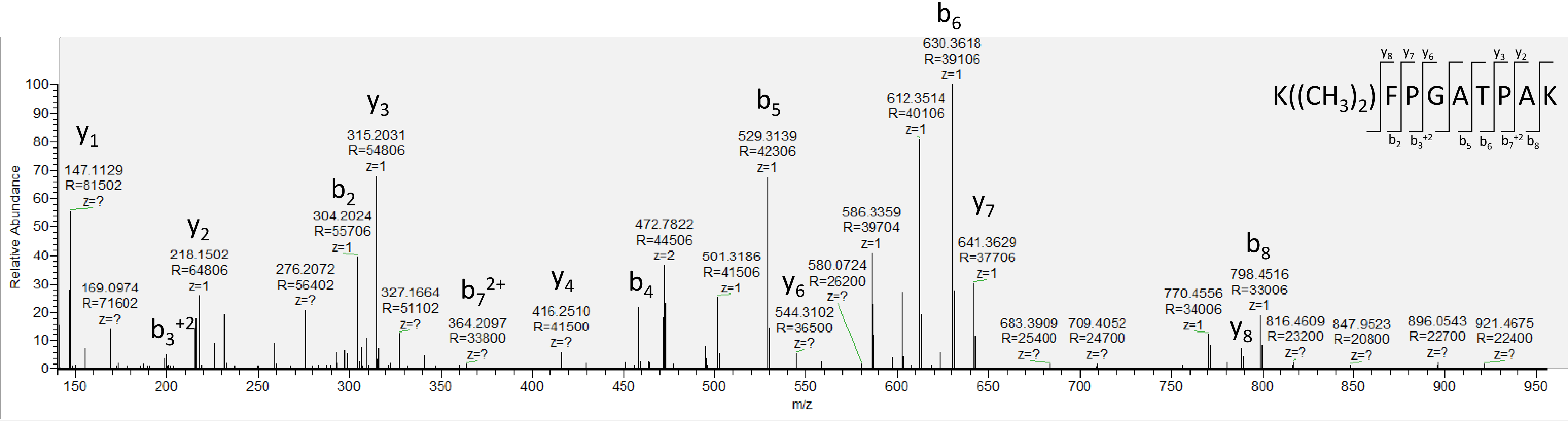

PALI1 K1241me2
Cell line: U2OS
PRIDE ID: PXD012354
(Ragazzini et al., 2019)

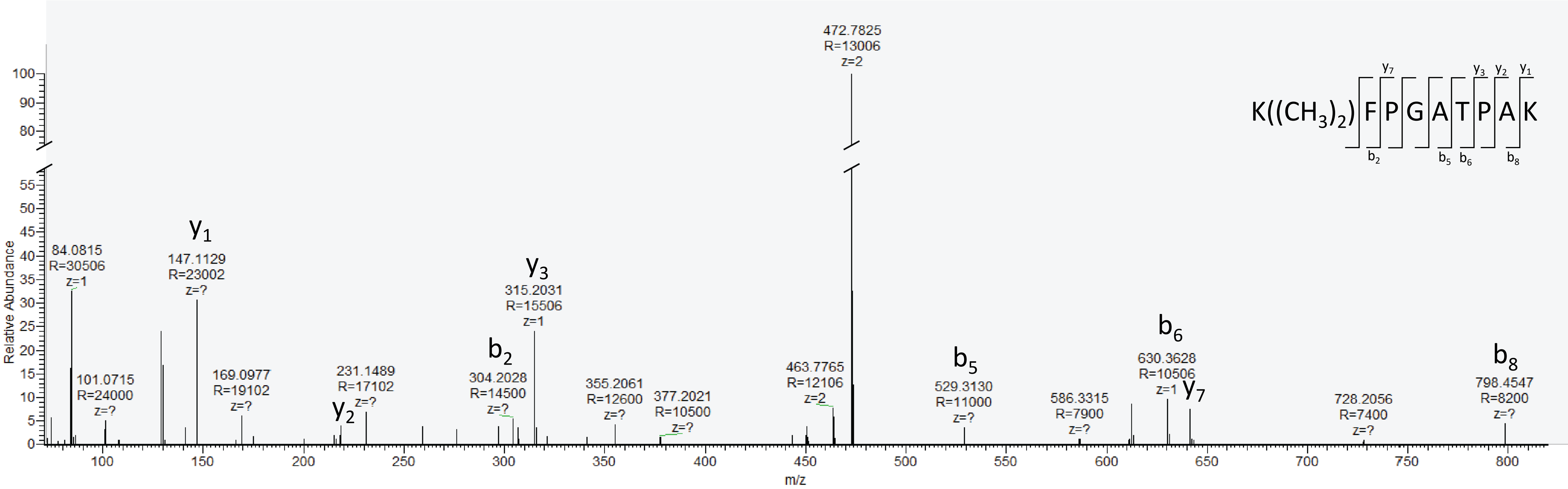

PALI1 K1241me2
Cell line: STS
PRIDE ID: PXD012547
(Wassef et al., 2019)

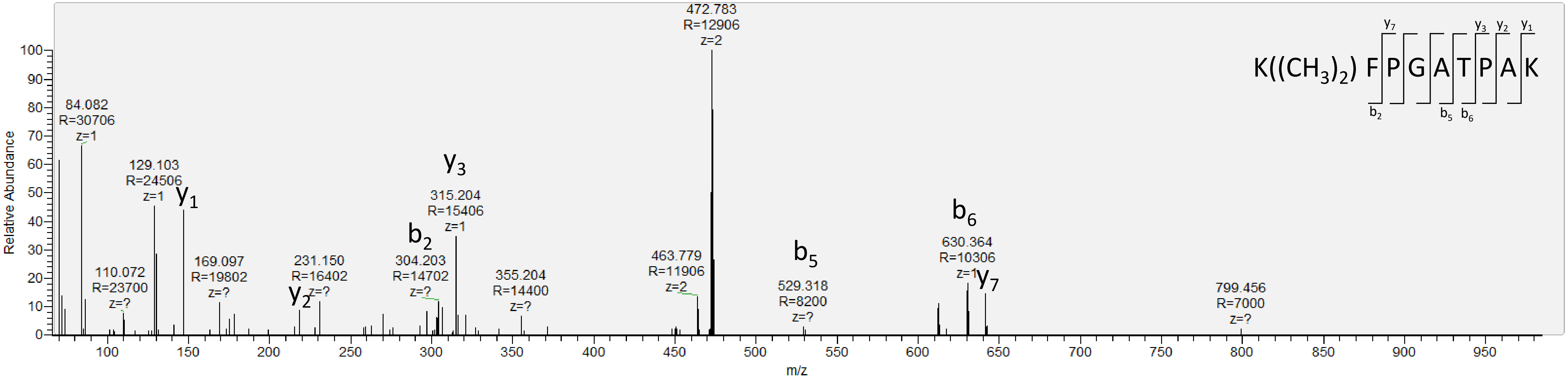

PALI1 K1241me2
Cell type: mESC
PRIDE ID: PXD003758
(Kloet et al., 2016)

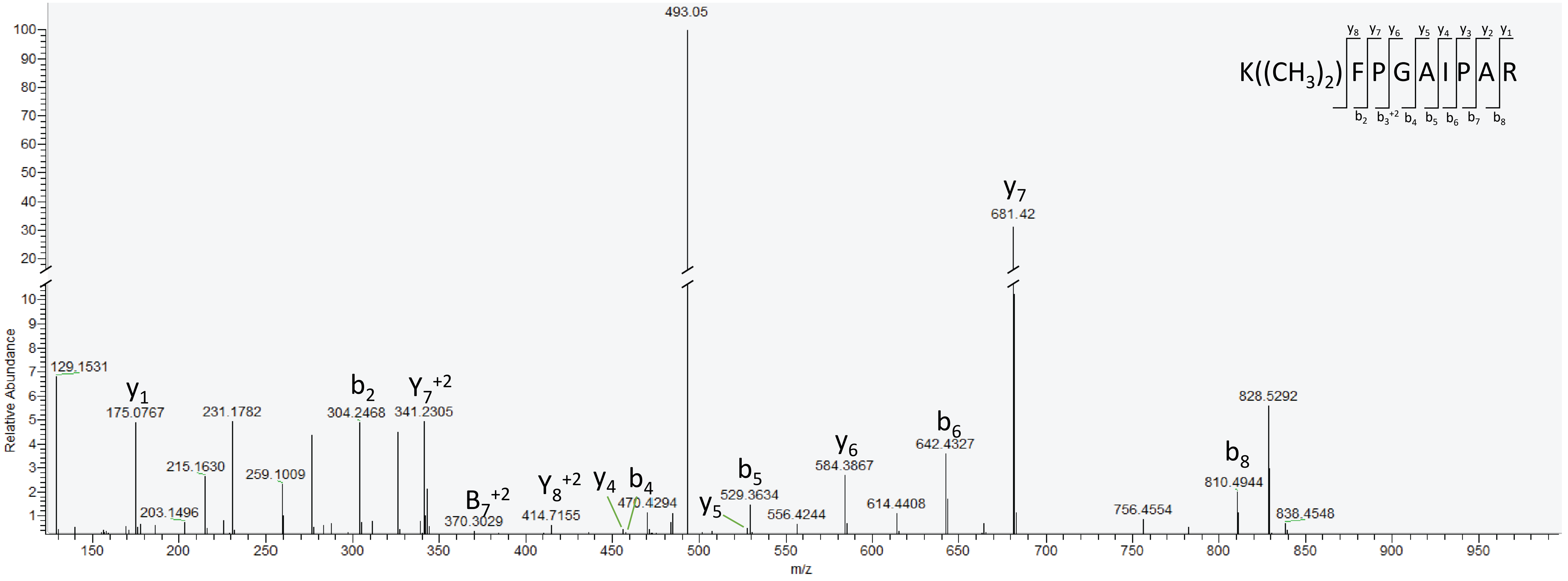

PALI1 K1241me3
Cell line: Ln-CAP
PRIDE ID: PXD012547
(Wassef et al., 2019)

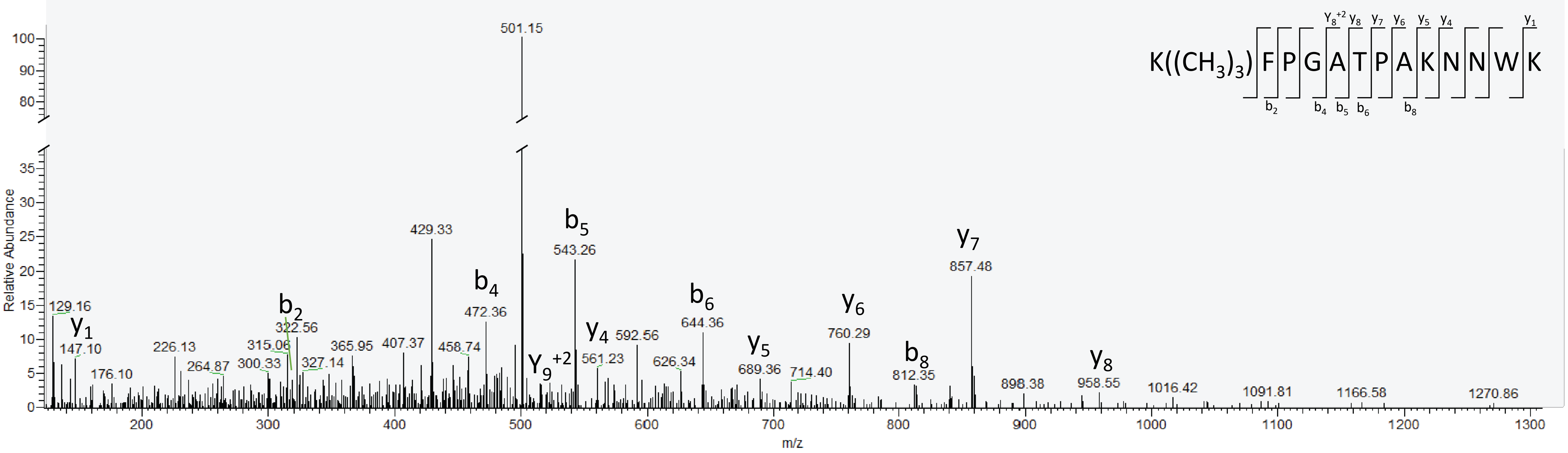
